# A cornea-specific role for the *Aspergillus fumigatus* carbon catabolite repressor, CreA, in tissue penetration and infection establishment

**DOI:** 10.64898/2026.08.13.744696

**Authors:** Becca L. Wells, Shi Ying Tang, Manali M. Kamath, Emily M. Adams, Jorge D. Lightfoot, Gautham S. Ramakrishnan, Can Zhao, Kevin K. Fuller

## Abstract

**Purpose:** Elucidate the influence of glucose metabolic pathways on *A. fumigatus* lung and corneal infection.

**Methods:** The *A. fumigatus acuF* and *creA* genes were deleted in an *mcherry*-expressing strain. The mutants were tested for alterations in radial growth, cell wall composition by fluorescence staining assays, and antifungal sensitivity through broth microdilution assays. Hyphal penetration of the strains through explanted porcine corneas was tracked by confocal microscopy using the mCherry signal. Virulence was evaluated in established models of invasive pulmonary aspergillosis (IPA) and fungal keratitis (FK) using C57BL/6J mice.

**Results:** Deletion of the *A. fumigatus* phosphoenolpyruvate carboxykinase (*acuF*) resulted in a dependency on exogenous glucose to support growth *in vitro*, but did not impact virulence in either the IPA or FK models. Loss of the carbon catabolite repressor CreA resulted in a broad dysregulation of carbon metabolic pathways and altered cell wall homeostasis. Surprisingly, whereas the *ΔcreA* remained fully virulent in the lung, the mutant was unable to establish infection in the FK model. This *in vivo* phenotype corresponded to an inability of *ΔcreA* to physically invade porcine corneal explants, which we attributed to a marked reduction in cell wall chitin content.

**Conclusions:** Gluconeogenesis is dispensable for *A. fumigatus* lung and corneal infection, suggesting tissue-derived glucose supports fungal growth in both environments. Loss of CreA disrupts glucose assimilation, its synthesis into chitin and, consequently, cell rigidity and hyphal invasion into the dense corneal stroma. Thus, CreA and other cell wall regulatory proteins may serve as targets for novel FK antifungals.

## 1. INTRODUCTION

*Aspergillus fumigatus* is a cosmopolitan fungus and predominant opportunistic pathogen. In patients with ablated or dysfunctional innate immunity, including bone marrow and organ transplant recipients, inhaled fungal spores (conidia) can germinate into hyphae that invade the lung parenchyma [1,2]. This disease entity, called invasive pulmonary aspergillosis (IPA), carries a mortality rate of 40-80% in treated cases and its incidence is expected to rise along with predisposing cancers and lung diseases such as COPD or COVID-19 [3,4].

*A. fumigatus* is also a common cause of fungal keratitis (FK), a corneal infection that impacts 1-2 million people annually [5]. The predominant risk factor for FK is not systemic immune suppression as in IPA, but rather damage to the otherwise protective corneal epithelium, most commonly due to ocular trauma or contact lens wear [6–8]. Once embedded into the cornea, *A. fumigatus* enjoys an immune privileged microenvironment ̶ marked by a low density of resident immune cells and an absence of infiltrating blood or lymphatic vessels ̶ which allows the fungus to establish infection before neutrophils infiltrate from the vascular periphery [9,10]. Though neutrophils play a critical role in fungal clearance, their presence refracts light (leading to opacification) and their secreted effectors drive stromal melt [11]. Contemporary antifungal therapies fail to resolve the infection in 30-60% of all FK cases, thus necessitating corneal transplantation or even enucleation if the infection spreads to the intraocular space [12].

Better antifungals are clearly needed to improve the survival or visual outcomes of *A. fumigatus* infections. Their development, however, requires the identification of putative drug targets, which we reason includes carbon assimilation pathways that support the energetic and biosynthetic demands of the fungus within the host tissue. In this study, we explore fungal carbon metabolism through the lens of one key metabolite: glucose. Glucose is central to fungal physiology, first as a structural building block for the cell wall polysaccharides (e.g., glucan and chitin) that define cell shape and provide resistance to various chemical or biomechanical stresses [13,14]. In carbohydrate-limiting environments, fungi must rely on *de novo* glucose synthesis (gluconeogenesis) using carbon derived from other substrates, such as acetate derived from the beta-oxidation of fatty acids [15,16].

Glucose is also the “preferred” energetic carbon source due to its relative ease of uptake through high-affinity transporters, coupled with its ATP-generating potential via substrate level and oxidative phosphorylation pathways. Fungi prioritize the utilization of glucose when it is available by repressing the expression of genes involved in alternative carbon (non-carbohydrate) utilization, e.g. ethanol or amino acid catabolism, thereby conserving the energy otherwise needed to drive those pathways [17,18]. This system, known as carbon catabolite repression (CCR), is governed by the *A. fumigatus* transcriptional regulator CreA [19–21]. Deletion of the *creA* gene in *A. fumigatus* strain CEA10 results in the up-regulation (de-repression) of roughly 400 genes, including those involved in ethanol utilization, the glyoxylate shunt (involved in acetate/two-carbon metabolism), and gluconeogenesis as expected. An almost equal number of genes are down-regulated in the mutant, suggesting CreA may function as both a transcriptional repressor and activator in *A. fumigatus* [19,20]. The result of this global metabolic dysregulation at any rate is a modest loss of fitness on all carbon sources, though the *ΔcreA* germination defect is most pronounced in glucose broth [19]. The mutant also displays an accentuated growth defect upon transition into hypoxia, which is attributable to reduced glucose fermentation, which supports growth when the respiratory capacity is reduced [19]. In a corticosteroid-induced mouse model of IPA, germination kinetics and the onset of animal mortality are similar between WT and *ΔcreA*-infected groups; however, the cumulative mortality at 14 days post-inoculation is approximately 30% lower for the mutant, which may reflect CreA’s role in optimizing glucose fermentation once hypoxia develops at later stages of disease [19]. To date, the influence of CreA or other glucose metabolic pathways on *A. fumigatus* pathogenesis in the cornea has not been evaluated.

In this study, we analyzed the virulence of gluconeogenesis and CCR deficient *A. fumigatus* mutants in murine models of IPA and FK. The data support a model in which the fungus assimilates glucose from the lung and corneal environments; however, only in the latter is the influence of CreA on glucose assimilation and cell wall homeostasis essential for tissue invasion and disease establishment. Taken together, these results demonstrate that the cornea presents unique stresses to the invading fungus, and that glucose metabolism or cell wall biosynthetic pathways may represent targets for novel FK therapeutics.

## 2. MATERIALS AND METHODS

### 2.1 Strains and culture conditions

All assays were conducted using minimal medium containing 1% w/v of the indicated carbon source, 10 mM ammonium tartrate, and salts (final pH=6.5) [21]. Knockout strains for *acuF* (Afu6g07720) and *creA* (Afu2g11780) were generated in an *A. fumigatus* reporter strain constitutively expressing mCherry (Af293, *PgpdA-mCherry-hph*) using established protocols [23–26]. Briefly, fungal protoplasts were incubated with: (1) Cas9 ribonucleoproteins (RNPs) designed to cut either side of the target coding sequence, (2) repair templates consisting of a bleomycin resistance cassette, from plasmid p402R, with 20 bp of homology flanking the cut sites, and (3) 60% polyethylene glycol in PBS. Transformed protoplasts were recovered on solid glucose minimal medium (GMM) supplemented with 1.2 M sorbitol and 125 μg/mL phleomycin. The reconstituted/complemented strains were generated by transforming the *ΔacuF* or *ΔcreA* protoplasts with the corresponding *acuF* or *creA* coding sequences, including 1,500 bp of the native promoter sequence (designed for ectopic integration). Protoplasts were co-transformed with *ptrA*, a pyrithiamine resistance gene that was amplified from plasmid p475. Transformants were recovered in 1 μg/mL pyrithiamine. Loss or reintegration of the target gene was confirmed via PCR of genomic DNA.

All oligonucleotides used in this study are listed in **Supplemental Table 1**.

### 2.2 *In vitro* growth assays

To evaluate sensitivity to various cell wall stressors or allyl alcohol, 2×10^3^ conidia were spotted onto glucose minimal media (GMM) supplemented with the various compounds as indicated. Plates were incubated at 35⁰C under atmospheric conditions. The colony diameter of each strain in the presence of the Congo red or calcofluor white was normalized to itself on GMM alone. For hypoxia phenotyping, inoculated plates were incubated at 35°C in normoxia (∼21% O_2_; 5% CO_2_) or hypoxia (1% O_2_; 5% CO_2_) for 72 h. Alternatively, plates were incubated in normoxia for 24 h and were either moved to hypoxia or allowed to continue growing under normoxic conditions for an additional 72 h incubation (96 h total). The ratio of colony diameter for each strain in hypoxia was normalized to itself in normoxia. Antifungal sensitivity was tested in a 96-well microdilution assay in GMM broth at 35⁰C. Micrographs were taken at 24 and 48h, and the optical density of each well at 530 nm was measured.

### 2.3 Cell wall polysaccharide profiling

GMM broth was inoculated with 10^4^ conidia/mL and incubated in a 6-well plate containing a coverslip in each well. Plates were incubated for variable times (16-20 h) at 35°C to ensure all strains reached the same development stage. Coverslips were washed 2x with PBS and hyphae were fixed with 4% paraformaldehyde for 15 min at room temperature. For aniline blue, coverslips were stained with 5 µg/mL aniline blue (pH 9.0) for 30 min in the dark at room temperature. Calcofluor white (fluorescence brightener 28) and alexafluor 350-conjugated wheat germ agglutinin (WGA, Invitrogen, W11263) staining was performed at room temperature using 10 and 5 µg/mL solutions for 15 min in the dark. In each case, excess stain was removed, and coverslips were washed once more with PBS before they were mounted with 90% glycerol on glass slides. Z-stack images (60X, 20-25 slices ea.) of hyphae were taken on an Olympus Fv1200 confocal microscope on the 60X setting using the DAPI (for aniline blue) and TxRed (for mCherry signal) channels. Fluorescence intensity was measured in average composite generated in ImageJ, and background fluorescence was subtracted to generate the corrected total cell fluorescence (CTCF). For each group, CTCF was measured in 9 hyphae across 3 biological replicates. Total β-glucan in the cell walls of WT, *ΔcreA*, and *ΔcreA* C’ hyphae was measured using a modified aniline blue protocol as described in the corresponding legend.

### 2.4 FK model

All FK studies were performed in accordance with the Association for Research in Vision and Ophthalmology (ARVO) guidelines for the use of animals in vision research and were approved by the University of Oklahoma Health Sciences Center Institutional Animal Care and Use Committee (Protocol: 20-060-CI). All procedures and analyses were performed as previously described [26–29]. Briefly, the corneas of methyl prednisolone-treated, 6-8 week old C57BL6/J male mice (Jackson Laboratories, Bar Harbor, ME, USA) were ulcerated and overlaid with metabolically activated (swollen) conidia. Contralateral eyes were sham-inoculated with PBS. Corneas were imaged daily through slit-lamp (Micron IV) or optical coherence tomography (Bioptigen) to assess clinical disease score and corneal thickness, respectively [26–29]. At 72 h post-inoculation, corneas were resected, digested in collagenase I (Millipore Sigma, SCR103), and plated onto inhibitory mold agar for CFU enumeration.

### 2.5 IPA model

Male 6-8 week old C57BL6/J (Jackson Laboratories) or CD-1 (Charles River Laboratories, Boston, MA, USA) mice received 40 mg/kg Kenalog-10 (triamcinolone) via subcutaneous injection at days −1, +2, and +5 relative to inoculation. On day 0, mice were inoculated intranasally with a 40 µL suspension of 5×10^7^ conidia/mL in PBS. Survival/moribundity was monitored for 14 days and percent survival was analyzed with a log-rank test. For fungal burden analysis, resected lungs were flash frozen in liquid nitrogen, lyophilized and homogenized with 0.1 mm zirconia/silica beads. Total DNA was extracted using the E.Z.N.A. ® Plant & Fungal DNA Kit (Omega Bio-Tek D3485). RNase-treated DNA was used for quantitative PCR, using TaqMan™ probes designed to amplify 18S region of *Aspergillus fumigatus* Af293. A standard curve of Af293 gDNA was run in parallel for linear regression analysis. For histology, lungs were fixed in 10% neutral buffered formalin and stored in 70% EtOH before staining with Grocott methenamine silver (GMS) or hematoxylin and eosin.

### 2.6 Quantitative RT-PCR

Strains were cultured as indicated in figure legends. RNA was DNase-treated using the DNase I kit (Millipore Sigma, Massachusetts, USA) and converted into cDNA using the ProtoScript II First Strand cDNA Synthesis Kit (New England Biolabs, Massachusetts, USA). The Luna Universal qPCR system (SYBR green; NEB, USA) was used to probe gene expression on the QuantStudio 3 Real-Time PCR System (Thermo Fisher Scientific, Massachusetts, USA). Fold-change in expression was calculated using the 2-ΔΔCt method.

### 2.7 NF-κB Activation Assay

To assess the antigenicity of the *ΔcreA* cell wall, RAW-Blue™ cells (Invivogen) were stimulated with heat-killed hyphal preparations of the various strains. NF-ĸB transcriptional induction was measured through secreted alkaline phosphatase. Details are provided in the corresponding figure legend.

### 2.8 *In vitro* penetration

A dual-layer agarose-based *in vitro* model was developed. The basal layer consisted of Hunter’s minimal medium solidified with 1% (w/v) agarose and poured into 90 mm petri dishes (25 mL per dish). Once solidified, 1 µL of spore suspension (containing approximately 100 conidia) was spotted, allowed to dry, and then overlaid with an agarose comprised of Hunter’s medium supplemented with either 3% or 5% (w/v) agarose (30 mL per plate). Plates were incubated at 37 °C for 72 h and then photographed.

### 2.9 *Ex vivo* lung penetration

To test the penetrative growth ability of the strains in the lung, 8-week-old male, immunocompetent C57BL6/J mice were intranasally inoculated with WT, Δ*creA*, or Δ*creA* C’ conidia (n=2/group, n=1 for PBS) as described above. After 1 h, mice were euthanized and their lungs were resected and plated whole onto petri dishes containing a thin layer of 1% agarose. Plates were imaged after incubation at 35⁰C for 48 h.

### 2.10 *Ex vivo* corneal penetration

Whole porcine eyes with clear, healthy corneas were obtained within 24 h post-mortem from a local abattoir (Manchester, UK). Corneas were excised at the limbus, leaving a 2–3 mm scleral rim; non-corneal tissues were removed. Corneal buttons were washed 3X in 5 mL sterile PBST-PS with gentle agitation and transferred, epithelium-side up, into individual wells of a 6-well culture plate containing 2 mL of PBST-PS medium. Each corneal button was superficially abraded with five parallel lines using a sterile 25G stainless steel needle (BD Microlance™, Fisher Scientific). Approximately 1 µL of spore suspension (∼10⁴ conidia/mL) was applied to the epithelial surface using sterile inoculation loops. Plates were left static at room temperature for 10–15 minutes to promote spore adherence, then incubated at 37 °C with 5% CO₂ for up to 48 hours. Uninfected controls received 1 µL of sterile PBS after epithelial scoring and were cultured under identical conditions. Culture medium was refreshed every 24 h by gentle pipetting.

For imaging, corneal buttons were placed epithelium-side down into 2-well ibidi imaging chambers (ibidi GmbH, Germany) containing 1 mL of the same culture medium used during infection. Live-cell confocal imaging was performed using a Leica TCS SP8 microscope (Leica Microsystems, UK) equipped with a 25× long working distance water-immersion objective. *A. fumigatus* expressing mCherry was imaged with excitation at 587 nm and emission collected between 600–630 nm. Fluorescence data were analyzed using Imaris v8.0 software (Bitplane, Switzerland), with fungal structures segmented using the ‘Surface’ module to assess tissue penetration depth. For each condition, three Z-stacks (1024 × 1024 µm, 300 µm thickness) were acquired per sample. The deepest observable point of fluorescent signal was recorded as the maximum penetration depth (µm); values exceeding 200 µm were capped at 200 µm.

### 2.11. Azocollagen hydrolysis assay

Collagenase secretion of *A*. *fumigatus* culture supernatants was quantified by Azocoll (194933; Millipore Sigma) hydrolysis as described previously [27,30] and in the corresponding figure legends.

### 2.12. Statistical analysis

Statistical analysis was performed on GraphPad Prism version 9.5.1. and the specific statistical tests applied for each experiment are described in the corresponding figure legends.

## 3. RESULTS

### 3.1. *A. fumigatus* gluconeogenesis is not required for establishment of IPA or FK in mice

To determine whether *A. fumigatus* relies upon gluconeogenic or exogenous (host-derived) glucose during infection, the gene encoding phosphoenolpyruvate carboxykinase, which catalyzes the first step in the gluconeogenic pathway [31,32], was deleted in an *mcherry* expressing reporter strain (Af293, *PgpdA-mcherry-hph*) [26]. Complementation of one such deletant (*ΔacuF*) was achieved through ectopic integration of the wild-type (WT) *acuF* allele (**Figure 1B &1C**). Growth of the isogenic set was first tested on media containing 1% w/v glucose and/or alternative carbon sources, including acetate, gelatin, and bovine serum albumin. In contrast to the WT and complemented (*ΔacuF* C’) strains that grew well on all carbon sources, *ΔacuF* grew only when glucose was supplemented (**Figure 1D)**. In acetate broth, the germination and hyphal extension rates of *ΔacuF* were improved with increasing glucose up to 250 µM, above which concentration the growth of the three strains was indistinguishable (**Figure S1A**). Notably, no additional phenotypes for *ΔacuF* under glucose-replete conditions were noted, e.g., altered sensitivity to oxidative or cell wall stresses (**Figure S1B**). We reasoned, therefore, that the pathogenic potential of *ΔacuF* would be tightly linked to the availability of glucose within the host milieu.

**Figure 1.**
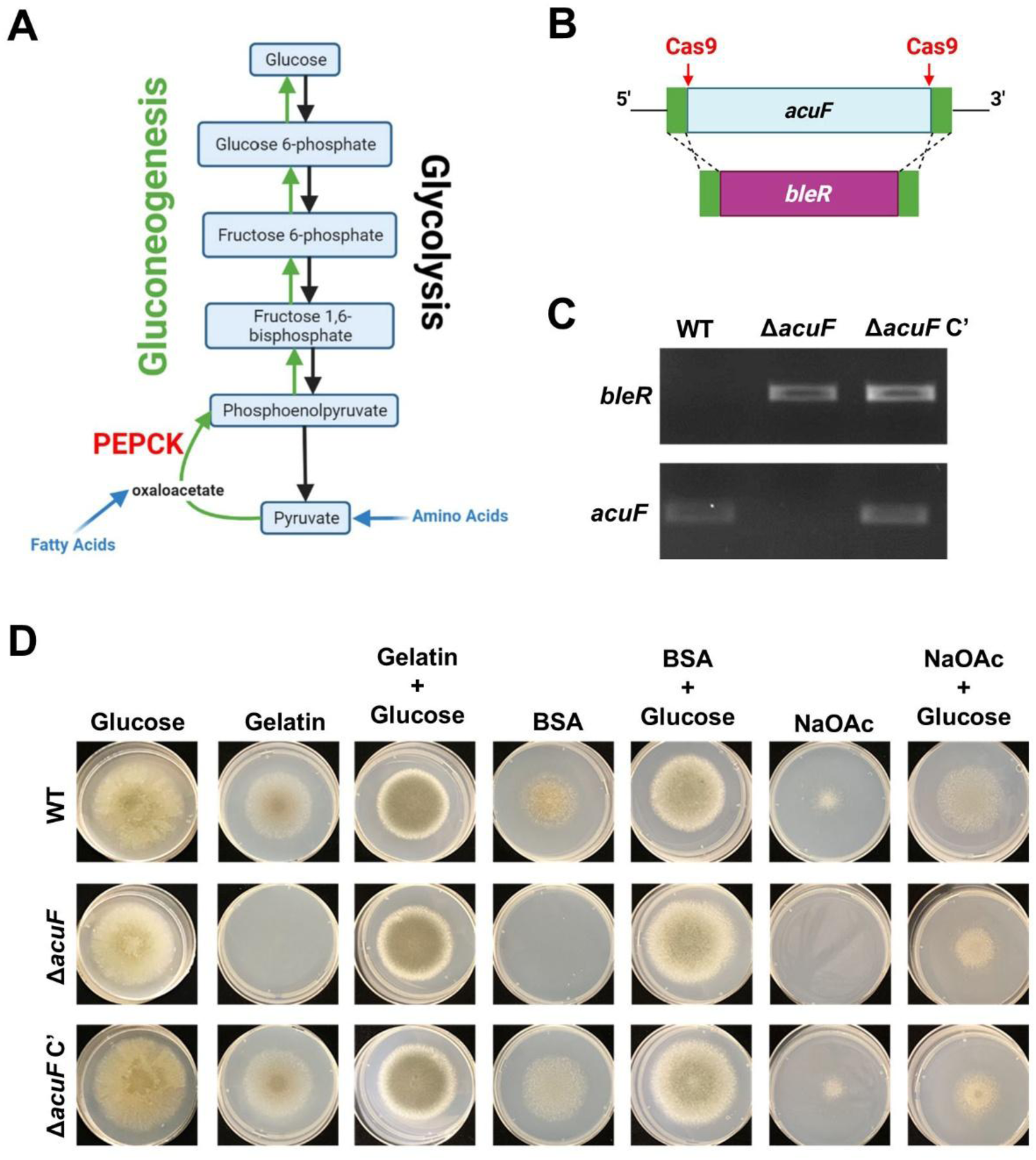
*A. fumigatus* gluconeogenesis (PEPCK) is required for growth in the absence of glucose. **A)** The enzyme PEPCK is required for the generation of glucose from citric acid cycle intermediates. Schematic was created using BioRender®. **B)** Cas9-mediated homologous recombination was used to replace *acuF*, the gene encoding PEPCK, with a bleomycin resistance cassette (*bleR*). **C)** The *acuF* deletant (Δ*acuF*) was confirmed via PCR of genomic DNA. **D)** The Δ*acuF* strain is unable to grow in minimal media containing complex carbon substrates (gelatin, bovine serum albumin, acetate) unless they are supplemented with 50 mM glucose. Images taken after 72 h incubation at 35°C.

Virulence of the *acuF* isogenic set was first tested in an established murine model of FK that involves ulceration of C57BL/6J mouse corneas followed by topical overlay with metabolically active conidia [26–29]. As shown in **Figure 2**, all strains drove comparable levels of clinical disease which corresponded to statistically indistinguishable fungal loads in corneas at 72 h. The strains were next evaluated in a model of IPA involving the intranasal instillation of fungal conidia into corticosteroid immunosuppressed C57BL/6J mice. At 48 h p.i., histological sections revealed similar degrees of inflammation and fungal outgrowth irrespective of the infecting strain (**Figure 2D**). This agreed with a separate cohort of animals in which statistically indistinguishable levels of fungal gDNA were detected between groups within total lung extracts (**Figure 2E**). The retention of virulence of the mutant was confirmed in a separate cumulative mortality study using outbred (CD-1) mice (**Figure S2**). Thus, *A. fumigatus* gluconeogenesis proved dispensable for virulence in the cornea and lung, suggesting that the fungus does not experience glucose starvation in either host environment.

**Figure 2.**
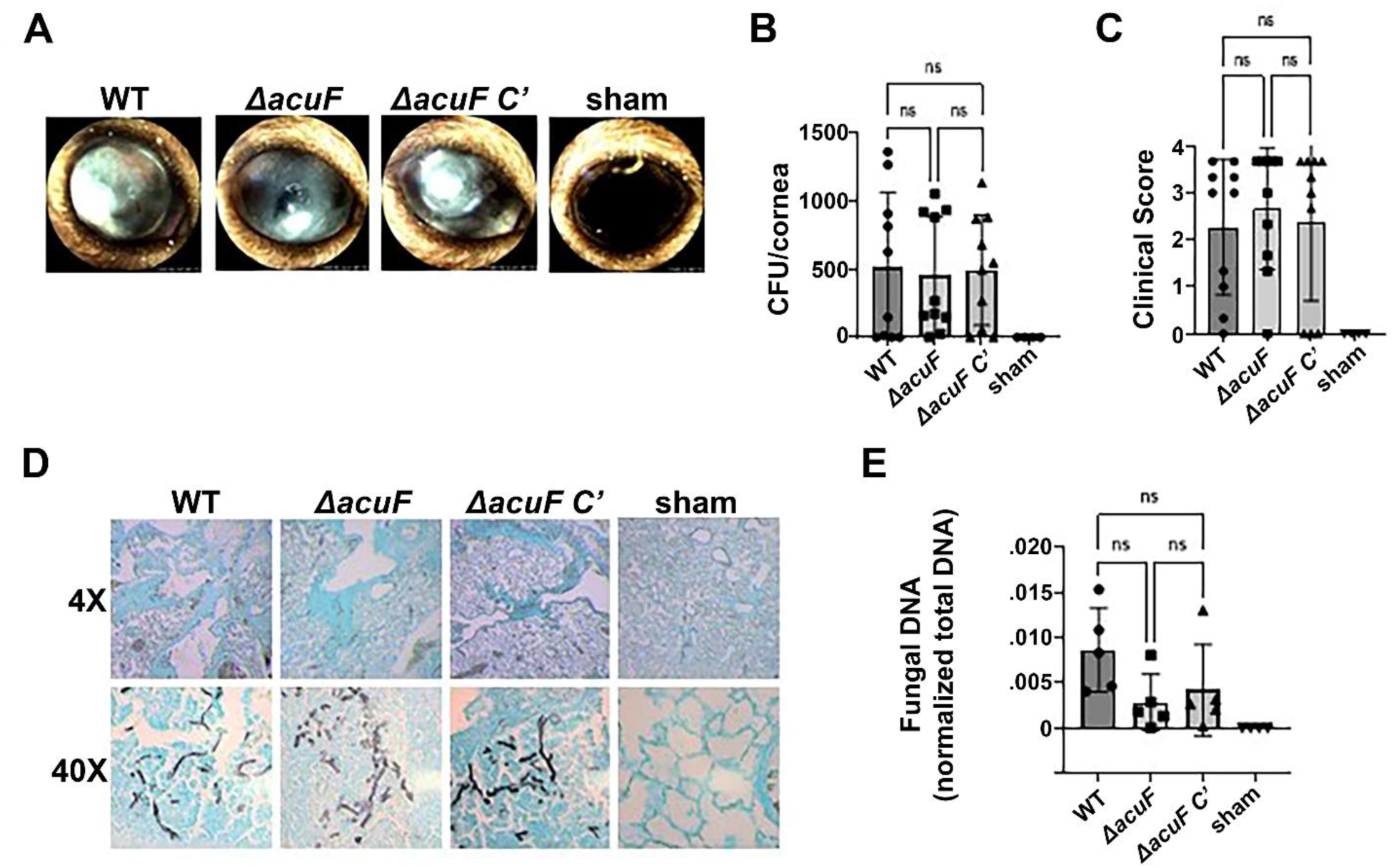
PEPCK is not required for *A. fumigatus* virulence in the cornea or lung. C57BL/6J corneas were ulcerated and topically overlayed with PBS (sham) or fungal conidia that were metabolically activated (swollen) in YPD medium (n=10/group, n=4 for sham). **A)** Representative slit-lamp images taken at 72 h post-inoculation (p.i.) reveal clinical manifestations of FK in all groups, while the sham-infected control appeared healthy. **B)** Slit-lamp images of infected corneas taken at 72 h p.i. were blindly scored 1-4 based on clinical severity. No differences in clinical score were observed between the groups (n=10/group, n=5 for uninfected). **C)** At 72 h p.i., corneas (n=10/group, n=5 for PBS) were removed, homogenized, and plated onto inhibitory mold agar (IMA). Colony forming units (CFU) were counted after overnight incubation at 35⁰C. (D-E) C57BL6/J mice were immunosuppressed with triamcinolone and intranasally inoculated with 2×10^6^ conidia (WT, Δ*acuF*, or *ΔacuF* C’) or PBS as a control. **D)** At 72 h post-inoculation (p.i.), lungs were removed (n=3/group, n=2 for PBS), sectioned, and stained Grocott’s Methenamine Silver (GMS). Whole scans of representative lungs are shown (40X magnification). **E)** DNA was isolated from lung homogenates (n=5/group, n=4 for Sham) taken at 72 h p.i. and fungal burden was measured via quantitative PCR of the *A. fumigatus* 18s rRNA gene. Panels B, C, and E were analyzed via Kruskal Wallis tests (ns=not significant).

3.2. Carbon catabolite repression regulates *A. fumigatus* metabolic fitness

The above results imply an important role for exogenous glucose assimilation during infection, which we next sought to disrupt by targeting CCR. We first generated *creA* deletion and complemented strains in our Af293 reporter background (**Figure 3A**) and evaluated CCR function of the strains in an allyl alcohol (AA) assay. Briefly, AA is metabolized into a toxic metabolite acrolein via enzymes encoded in the alcohol dehydrogenase-aldehyde dehydrogenase (*alcA*/*aldA*) gene cluster, which is repressed by CreA in the presence of glucose [19]. Consequently, and in accordance with our results, both the WT and complemented strains grew in the presence of 0.1% AA on glucose (repressing) media but were inhibited by AA on de-repressing gelatin (collagen lysate). By contrast, *ΔcreA* was inhibited by AA irrespective of the carbon source, suggesting that the *alcA* cluster was constitutively de-repressed and CCR thus abolished in the mutant (**Figure 3B**). In the absence of AA, the colony diameter of the *ΔcreA* on either gelatin or glucose was indistinguishable from the WT at 24 h p.i., suggesting that CreA does not influence germination/colony establishment. The linear expansion rate between 24 and 72 h was however reduced in the *ΔcreA* mutant on both media, where this defect was more pronounced on glucose (**Figure 3C**). Since *A. fumigatus* shifts its metabolism towards glucose metabolism (fermentation) in low oxygen conditions [33], we next tested whether the glucose metabolic defect of *ΔcreA* impacted hypoxic growth. As shown in **Figure 3D**, the growth of *ΔcreA* in 1% O_2_ was not markedly altered relative to itself in normoxia. Similarly, the sensitivity of the *creA* isogenic set to various oxidative stressors, including hydrogen peroxide, paraquat, and tert-butyl alcohol, was similar (**Figure S5**).

**Figure 3.**
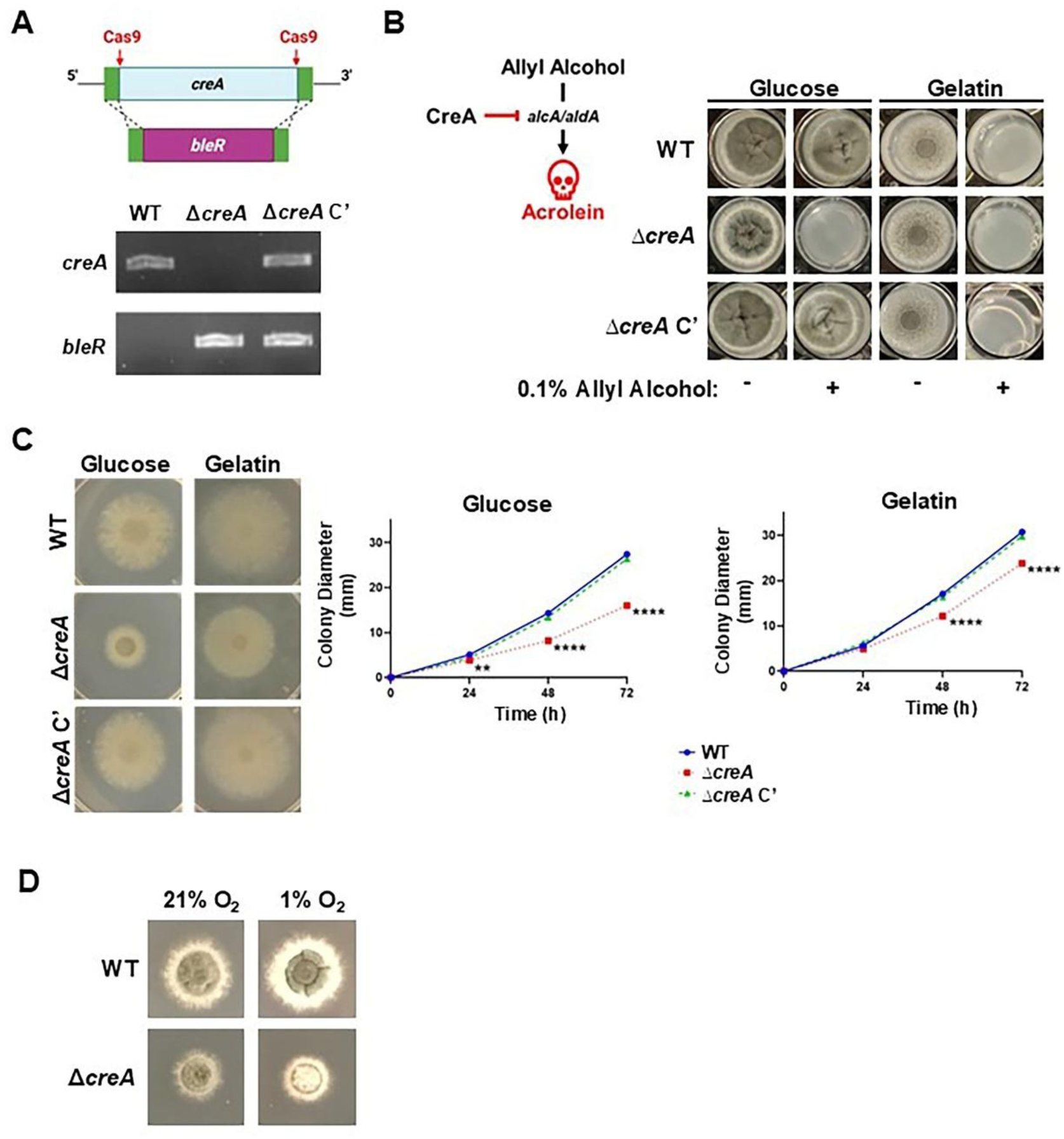
Loss of CCR decreases *A. fumigatus* fitness under repressive and de-repressive conditions. **A)** The *creA* coding sequence was deleted and replaced with a bleomycin resistance cassette in an *A. fumigatus* Af293 background. Deletion was confirmed via PCR of genomic DNA. Schematic created with BioRender ®. **B)** The inability to grow in the presence of allyl alcohol, which is metabolized into the toxic product acrolein by genes encoded in the CCR-respondent *alcA*/*aldA* gene cluster, phenotypically confirmed loss of CreA function in the Δ*creA* strain. **C)** Conidia were spotted onto minimal medium containing glucose or gelatin as the sole carbon source. Images were taken after 72 h incubation at 35⁰C. Radial growth diameter of each strain was measured at 24, 48, and 72 h and analyzed via One-way ANOVA (**P<0.01, ****P<0.0001). **D)** Strains were spotted in triplicate onto glucose minimal medium and incubated at 35⁰C in normoxia (atmospheric oxygen; 19-21%) or in low oxygen (1% O_2_) for 48 h. Representative images shown.

### 3.3 CreA influences the cell wall polysaccharide composition and hyphal morphology

Given that glucose serves as an anabolic substrate for cell wall polysaccharides [13], we next evaluated the influence of CCR disruption on cell wall homeostasis. To begin, the sensitivity of the *creA* isogenic set was tested against Congo red and calcofluor white, which drive cell wall stress by binding and inhibiting glucan and chitin polysaccharides, respectively [34,35]. As shown in **Figure 4**, the supplementation of either compound into glucose medium reduced the colony diameter of the Δ*creA* mutant (relative to itself on control medium) to a significantly greater extent than either the WT or complemented strains. The Δ*creA* mutant was also hypersensitive to the antifungal caspofungin, which inhibits the activity of the β-glucan synthase enzyme [36,37] (**Figure 4C**).

**Figure 4.**
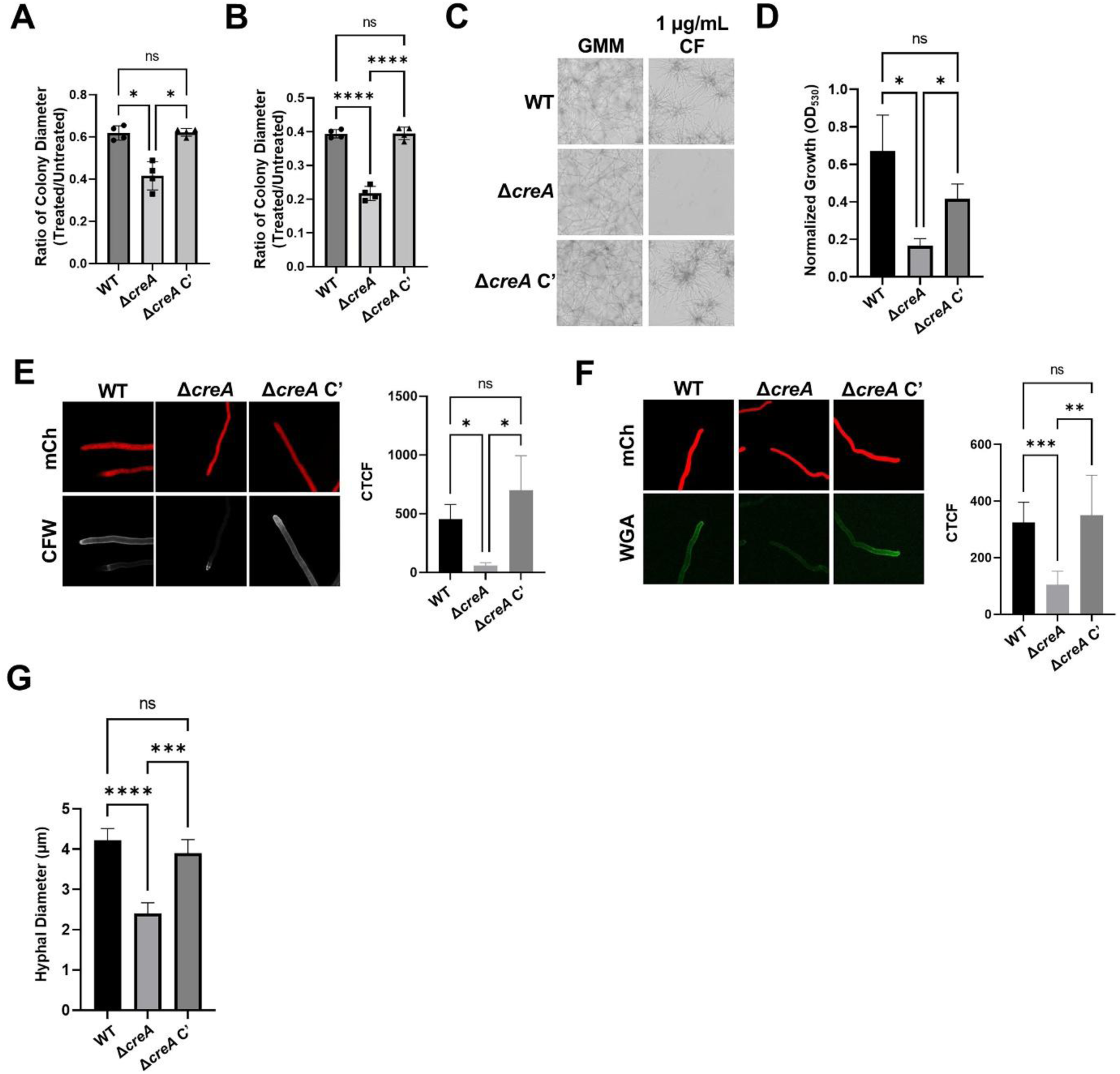
**Cell wall composition is altered in the Δ*creA* strain**. Conidia of WT, Δ*creA*, and Δ*creA* C’ were spotted onto GMM alone (untreated) or supplemented with 20 µg/mL Congo red **(A)** or 15 µg/mL calcofluor white **(B)**. The colony growth diameter was measured after 72 h incubation at 35⁰C and the growth of each strain in the presence of the cell wall stressors was normalized to its average colony growth diameter on untreated media (n=4/group). Analyzed via Brown-Forsythe and Welch ANOVA test (*P<0.05, ****P<0.0001). **C)** 5×10^4^ conidia of each strain were seeded into the wells of a 96-well plate containing GMM alone or with 1 µg/mL caspofungin and incubated at 35⁰C. Representative micrographs were taken at 48 h. **D)** The optical density (OD) of the wells at 530nm was measured and the OD of each strain in caspofungin was normalized to its baseline OD in GMM (n=4/group). **E-G)** Strains were grown on sterile coverslips at 35⁰C in GMM for 18-24 h before they were stained with calcofluor white **(E)** to visualize total chitin in the cell wall or FITC-conjugated wheat germ agglutinin **(F)** to determine levels of chitin exposed at the hyphal surface. Composite images were generated from z-stack images taken on an Olympus FV1200 confocal microscope and corrected total cell fluorescence (CTCF) was quantified in ImageJ and analyzed via Brown-Forsythe and Welch ANOVA tests (n=9-10/group). **G)** The hyphal diameter was measured for each of the strains using the Mcherry signal (n=5/group; groups were analyzed via Ordinary One-way ANOVA (*P<0.05, **P<0.01,***P<0.001, ****P<0.0001).

The cell wall polysaccharide content of the *creA* strains was assessed by staining glucose-grown hyphae for β-glucan with aniline blue and soluble dectin-1 receptor, or chitin with calcofluor white and fluorophore-conjugated wheat germ agglutinin [38–41]. Interestingly, morphometric analysis of hyphal images across the various stains revealed a consistent and statistically relevant reduction in hyphal diameter for the *ΔcreA* mutant (**Figure 4G**). Nevertheless, and as shown in **Figure S3**, the aniline blue and dectin-1 staining signals were statistically indistinguishable across the strains, suggesting β-glucan content was similar between the strains. By contrast, the intensity of both chitin stains was markedly reduced in Δ*creA* (**Figure 4E and 4F).** To determine whether reduced chitin could be attributed to a transcriptional influence of CreA on chitin wall related genes, the expression of several chitin synthases was subsequently evaluated under the same culture conditions. As shown in **Figure S3D**, *chsA* displayed a modest (2-fold) reduction in expression in Δ*creA*, whereas the expression of others, including *chsB* and *chsE*, was unaltered.

Fungal cell wall polysaccharides serve as the principal antigens recognized by innate immune cells in the lung and cornea [42,43], suggesting that loss of CreA may influence the host-fungal interaction. However, heat-inactivated hyphae of WT and *ΔcreA* drove comparable levels of NF-ĸB activation in RAW macrophages, suggesting that the hyphal cell wall antigenicity was not altered in the mutant (**Figure S3C**).

### 3.4 CreA regulates infection establishment in a niche-dependent manner

The virulence of the *creA* isogenic set was first tested in the FK model described above. Remarkably, and in contrast to the progressive disease course observed in corneas infected with the WT or complemented strains, those inoculated with the *ΔcreA* failed to display clinical signs of disease upon slit-lamp imaging or changes in corneal thickness (edema) based on optical coherence tomography (**Figure 5A and 5B**). The absence of disease in *ΔcreA* corneas corresponded to a lack of viable fungus within the tissue at 72 h p.i., indicating that the mutant was incapable of establishing infection in this model (**Figure 5D**).

**Figure 5.**
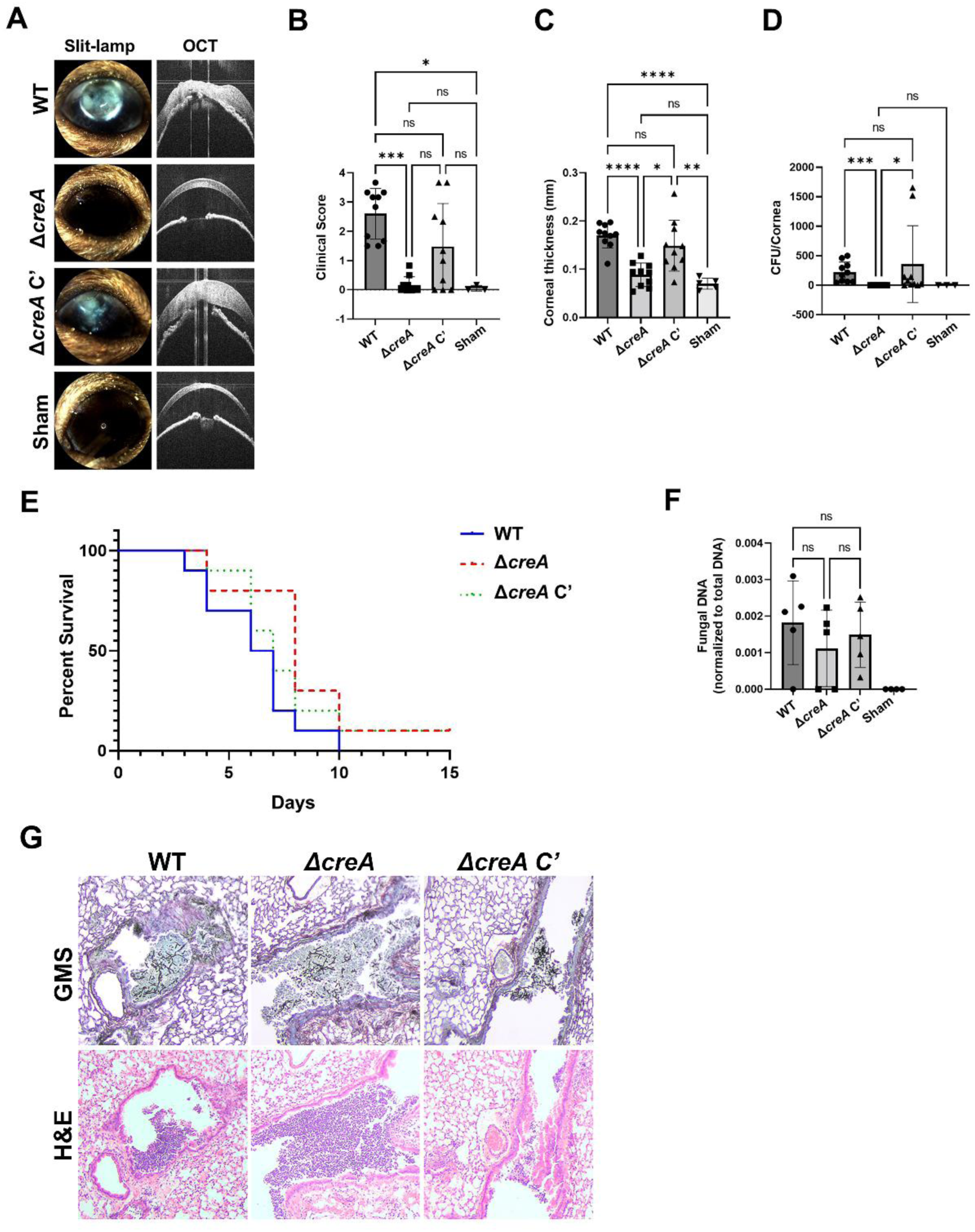
CreA is essential for the establishment of infection in the cornea, but not the lung. The corneas of male C57BL/6J mice were ulcerated and topically inoculated with WT, Δ*creA*, or *creA* C’ conidia and infections were allowed to develop for 72h. **A)** Representative slit-lamp and OCT images taken at 72h demonstrate clinical signs of infection in the WT and *creA* C’ inoculated mice, but not the Δ*creA* inoculated mouse. **B)** Slit-lamp images of infected corneas taken at 72h p.i. were blindly scored 1-4 based on clinical severity. Disease scores were significantly reduced in the Δ*creA* group compared to the WT and *creA* C’. **C)** OCT was used to measure corneal thickness at 72h p.i., which was also significantly reduced for Δ*creA* inoculated mice. **D)** After 72h, corneas were collected, homogenized, and plated onto IMA. After overnight incubation at 35⁰C, colony forming units (CFU) were counted. Groups in panels **B-D** were compared via Kruskal Wallis tests (*P<0.05, **P<0.01,***P<0.001, ****P<0.0001; n=10/group, n=3 for Sham). For **E-G**, C57BL/6J mice were immunosuppressed with triamcinolone and inoculated intranasally with 2×10^6^ conidia. **E)** The survival curves for each infection groups were indistinguishable by Logrank test (n=12/group); all groups were statistically different from PBS (sham) inoculated controls. **F)** Fungal burden in lungs taken from mice inoculated as described above at 48h p.i., measured by qPCR of the Af293 18S rDNA region (n=4-5/group) and analyzed via Kruskal Wallis tests (*P<0.05, **P<0.01,***P<0.001, ****P<0.0001). **G)** Lungs taken at 48 h p.i. reveal comparable levels of fungal growth within the bronchiolar space upon Gamori Methenamine Silver (GMS) staining in which fungal hyphae appear black.

The virulence of the strains was next tested in the steroid model of IPA using C57BL/6J mice. In a cumulative mortality study, animals began to succumb to infection at 4 or 5 days p.i. irrespective of the infecting strain, and the two-week survival curves were statistically indistinguishable between the infected groups (**Figure 5E**). In an independent experiment, lungs of inoculated mice were harvested at 72 h p.i. and showed comparable fungal loads based on qPCR analysis as well as comparably sized fungal and inflammatory lesions in histological sections (**Figure 5F and 5G**). Taken together, the influence of CreA on *A. fumigatus* virulence varies markedly between the FK and IPA models.

### 3.5 The Δ*creA* virulence correlates with its penetration phenotypes in corneal and lung tissue

Whereas the lung parenchyma is defined by a spongy (open-air) interface that facilitates gas exchange, the corneal stroma is markedly more dense (semi-crystalline) due to the compact deposition of collagen fibers needed to achieve optical clarity [44,45]. Since the *ΔcreA* wall is deficient in chitin ̶ which regulates cell wall rigidity and maintains morphology under biomechanical stress [46,47]– we hypothesized that the virulence defect of the mutant in the cornea is tied to an inability to physically penetrate the stromal matrix. To test this, we first assessed the impact of substrate density on fungal growth. Conidia of the WT, Δ*creA*, and *creA* C’ strains were spotted onto complete medium containing 1% agarose, subsequently overlaid with 3% or 5% agarose medium, and incubated at 37°C for 48 h. As shown in **Figure 6A**, all three strains were able to penetrate through the 3% medium comparably well, whereas the *ΔcreA* mutant was notably more impaired at 5%.

**Figure 6.**
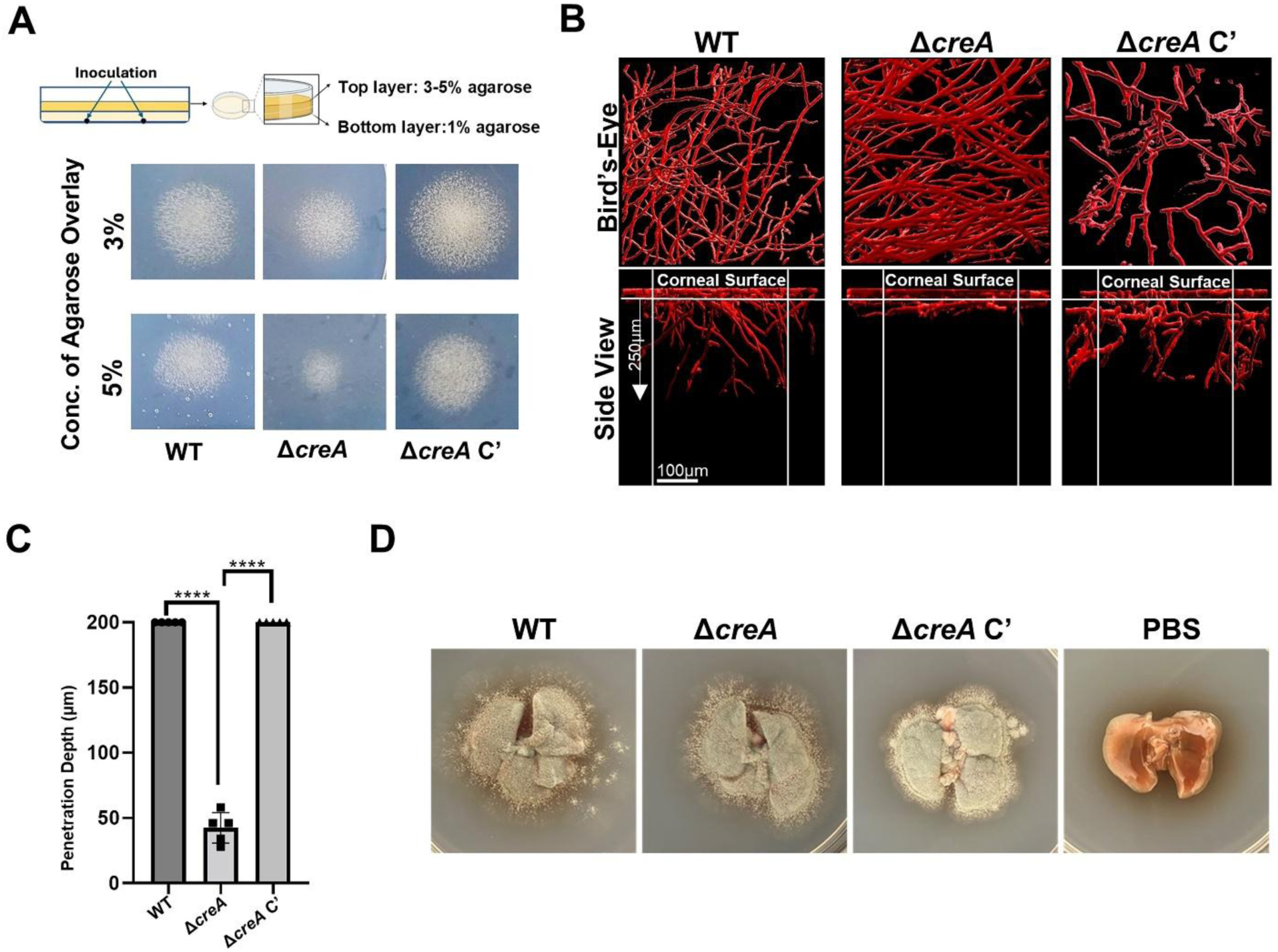
**The Δ*creA* strain displays a penetrative growth defect in *ex vivo* corneas, but not lungs**. **A)** WT, Δ*creA* and Δ*creA* C’ were spotted onto minimal media containing 1% agarose and dried spots were subsequently overlaid with media containing 3 or 5% agarose then incubated for 72h at 37⁰C to determine the ability of each strain to grow in a penetrative manner. **B)** *Ex vivo* porcine corneas were placed into 6 well plates, manually abraded at the surface, and topically inoculated with WT, Δ*creA* or Δ*creA* C’. Plates were incubated at 37⁰C for 48h and fungal growth across the surface (Topical) and down into the deeper layers (Vertical) of the cornea was visualized on a Leica TCS SP8 live-cell confocal microscope (excit: 587 nm, em: 600–630 nm). The Δ*creA* strain was unable to penetrate down into the corneal matrices. **C)** Z-stack images were analyzed using Imaris v8.0 software and used to quantify the depth to which the fungal growth penetrated into the corneal tissue (capped at 200 µm). The Δ*creA* strain’s growth was confined primarily to the corneal surface (≤50µm). Groups were compared by Ordinary One-way ANOVA (****P<0.0001). **D)** C57BL/6J mice were inoculated with 2×10^6^ conidia intranasally (n=2/group) and whole lungs were collected and plated onto 1% agarose after 1hr, then incubated at 35⁰C. All strains were able to penetrate the lung tissue and grow through to the surface after 48h.

To determine if the density-dependent growth defect of *ΔcreA in vitro* correlated with corneal invasion, resected porcine corneas were inoculated topically with the *creA* isogenic set and incubated at 37⁰C for 48 h. The cytoplasmic mCherry signal from the strains was then used to track and quanitfy fungal biomass on and within the tissue by confocal microscopy. As shown in **Figure 6B**, the *en face* (“bird’s eye”) imaging revealed that hyphal accumulation on the corneal surface was indistinguishable across the three strains. Strikingly, however, the cross-sectional (“side view”) images revealed that whereas the WT and complemented strains had penetrated the stromal matrix to comparable depths, hyphae of the *ΔcreA* mutant were restricted to the inoculated surface (**Figure 6B and 6C**).

To test penetration of the strains through lung tissue, immunocompetent C57BL/6J mice were inoculated intranasally with 2×10^6^ conidia and immediately resected and plated onto agarose medium for incubation at 35°C. As shown in **Figure 6D**, each of the strains penetrated through to the lung surface within 48 h (Figure 5D), indicating that loss of CreA did not impair penetrative growth through the lung.

## 4. DISCUSSION

As glucose is an essential nutrient or metabolite for fungal growth, it follows that the disruption of glucose homeostasis might serve as an effective antifungal modality. Critically lacking, however, has been an understanding of which glucose metabolic pathways are operative during infection (e.g. catabolic versus anabolic) or if such programming might differ as a function of host niche. Our analysis of a gluconeogenesis-deficient *A. fumigatus* mutant, a first for this fungus, begins to address these questions. Growth of *ΔacuF in vitro* was dependent upon an exogenous supply of glucose as predicted and became indistinguishable from the WT at concentrations above 0.25 mM. The capacity of *ΔacuF* to establish infection in the FK and IPA models strongly indicates, therefore, that the bioavailability of glucose is above this threshold in both tissues. This interpretation is compatible with more formal estimates. The glucose concentration of lung airway surface liquid, for example, has been measured at 0.4 mM [48,49], which despite being 3-20X lower than that found in plasma, should nevertheless support growth of the *ΔacuF* mutant. This surplus of glucose in the lung, at least from the fungus’ perspective, may explain why mutants defective in alternative carbon assimilation pathways ̶ including the two-carbon, lipid, or protein metabolism– retain their virulence within murine models of IPA [**5**0-52]. Glucose levels in the cornea have been modeled to be even higher (ranging from 1-7 mM), the main source of which being the aqueous humor produced in the anterior chamber [53,54]. Taken together, the data from this and prior studies support a model in which *A. fumigatus* assimilates adequate levels of glucose in both the lung and cornea to support its energy and cell wall synthesis demands.

We next reasoned that shifting the metabolic state of *A. fumigatus* away from exogenous glucose use, achieved here through the ablation of CCR, would attenuate virulence in both the IPA and FK models. We were therefore surprised to find the *ΔcreA* mutant was fully virulent in lung but avirulent in the cornea, leaving us to consider key differences between the two tissues that might account for this. The loss of CreA results in a dysregulation of roughly 10-20% of the *A. fumigatus* genome on both repressing and de-repressing carbon sources *in vitro*, where most of the affected genes are related to primary and secondary metabolism [19,20]. The *ΔcreA* mutant is consequently maladapted and slower-growing than WT under all nutrient conditions, though this defect is accentuated in glucose-replete environments. Nevertheless, the mutant retained its ability to grow on the corneal surface *ex vivo*, suggesting the nutritional acquisition *per se* is not the principal barrier to *ΔcreA* infection in the FK model. Additional parameters that are unique to the corneal microenvironment thus need to be considered.

Oxygen is a key environmental factor that shapes carbon metabolism in obligate aerobes such as fungi. Regarding infection, it has been shown that *A. fumigatus* shifts its metabolism towards glucose fermentation in the murine lung as necrosis and infiltrating leukocytes drive tissue hypoxia [33]. Thus, while the glucose assimilation defect of *ΔcreA* appears not to impact infection establishment in the healthy airway (i.e., where other carbon sources could in principle be oxidized or shunted into gluconeogenesis), it may affect fitness and persistence at the later, oxygen-depleted stages of disease. Indeed, Beattie et al., in the *A. fumigatus* CEA10 background, observed reduced mortality in *ΔcreA*-inoculated animals in the latter half of a two-week experiment ̶ a result, it should be noted, that was not recapitulated in the present study, which differs in both fungal and mouse background [19]. Nevertheless, the oxygen dynamics during FK are different, as we have shown that *A. fumigatus* infected corneas become hypoxic as early as 6 h post-inoculation [26]. This precedes clinical signs of corneal inflammation and may result instead from an increased oxygen consumption by antigen-stimulated corneal resident cells. This rapid onset of hypoxia may drive a heightened dependency of *A. fumigatus* on glucose fermentation at the early stages of FK, relative to the IPA, and account for the avirulent *ΔcreA* phenotype in the cornea. Though this cannot be ruled out, the lack of a robust hypoxic growth phenotype for *ΔcreA in vitro* does indicate other environmental parameters are likely more salient. Tissue density, for example, also varies markedly between the lung and cornea: a spongy lung parenchyma to support gas exchange versus a semi-crystalline (highly dense) corneal stroma that serves as an optical lens [44,45]. We reason that this mechanical difference underpins the penetration phenotype of *ΔcreA* through explanted tissue, which in turn is predictive of its virulence in the FK and IPA models.

*A. fumigatus* penetration through stromal collagen may be facilitated in principle through the secretion of extracellular proteases, which account for approximately 1% of the entire genome. Indeed, we previously demonstrated that several protease-encoding genes are upregulated 10-1000 fold in the murine cornea (relative to standard culture conditions), though the environmental signal(s) that trigger the transcriptional response remain unclear [27]. We argue, however, that a deficiency in proteolytic activity does not account for the tissue invasion or virulence defect of *ΔcreA*. First, we did not see a reduction in collagenolytic activity in *ΔcreA* conditioned media and (**Figure S5**), second, we previously demonstrated that a marked ablation of protease activity in *A. fumigatus* (via deletion of protease-specific transcription factors) does not impact virulence in the FK model [27].

Another determinant of fungal invasiveness is the cell wall, which must be rigid enough to counter the biomechanical stress applied by the external substrate. This rigidity is loosely proportional to the level of cell wall chitin, a linear polymer of β-1,4-linked N-acetylglucosamine (GlcNAc) with a tensile strength exceeding bone and steel [46]. Chitin’s structural strength can indeed be appreciated outside of the Fungi, as it is the principal component of the arthropod exoskeleton [55]. Our working hypothesis, therefore, is that the marked reduction in total chitin observed in the *ΔcreA* wall critically impacts penetration through the corneal stroma, but not the lower density lung. Given the pleiotropic nature of CreA, this model would be best supported through an independent analysis of a chitin-specific mutant. This is complicated however by the fact that *A. fumigatus* encodes eight highly redundant or compensatory chitin synthases [56]. A discernable (70%) reduction in mycelial chitin content has only been observed in a quadruple knockout (*ΔchsA/csmB/chsF/chsG*) that is profoundly growth inhibited and does not sporulate, thus rendering it incompatible for virulence testing [57]. We reason that the growth and developmental phenotypes of *ΔcreA* reflect a more modest reduction in cell wall chitin, but one that is nevertheless sufficient to impact invasive growth. On this topic, it is interesting to note that a different quadruple chitin synthase mutant (*ΔchsE/Eb/D/F*), despite not having an appreciable reduction in cell wall chitin at baseline, is hypersensitive to Congo red and caspofungin (like *ΔcreA*) as well as neutrophil-derived chitinase (AMCase) treatment *in vitro*. C57BL/6 corneas inoculated with this mutant show less clinical disease and harbor less fungus than WT-infected controls, and this phenotype is lost in an AMCase^-/-^ mouse background [58]. Though this work with the *ΔchsE/Eb/D/F* mutant cannot speak to the impact of a baseline chitin defect like the one seen in *ΔcreA*, it does suggest that altered fungal-neutrophil interactions may play a role in the *ΔcreA* virulence phenotype. This is currently under investigation in our group.

The mechanism through which CreA influences cell wall chitin in *A. fumigatus* also remains incompletely understood. It is well described in this and other fungi that the chitin synthases, which polymerize UDP-N-GlcNAc-1-phosphate monomers, are regulated by transcription factors such as RlmA downstream of the PKC-MAPK cell wall integrity (CWI) pathway [59–62]. It follows that CreA, in its capacity as a transcriptional regulator, may regulate chitin synthase expression as part of, or in parallel to, canonical CWI signaling. However, prior studies in which the *ΔcreA* transcriptome was analyzed did not reveal significant alterations CWI kinase expression, *rlmA*, or any of the eight chitin synthase genes on glucose medium [19,20]. Though our qRT-PCR results do suggest a modest (2-fold) reduction in the *chsA* and *chsE* genes in *ΔcreA*, this likely does not fully account for the marked chitin reduction given the functional redundancy across the chitin synthase paralogs. Rather, we hypothesize that the global dysregulation of carbon metabolism in *ΔcreA* impacts the flux of glucose towards chitin biosynthesis, which includes glucose uptake and phosphorylation, isomerization to fructose, amination to glucosamine, and acetylation to form the GlcNAc precursors. Interestingly, we did not appreciate a comparable reduction in total β-glucan, which is a glucose polymer, the predominant cell wall polysaccharide and principle fungal antigen. Beyond the cell wall staining analyses, we observed that the WT and *ΔcreA* strains also stimulated NF-ĸB expression cultured macrophages to a similar extent, further suggesting that glucan levels and exposure are not altered in the mutant. More work is needed to fully understand the fate of glucose across the various polysaccharide pathways in the WT vs *ΔcreA* strains.

Though critical for the establishment of FK, it is currently unclear to what extent CreA or chitin dynamics influence pathogenesis at later stages of disease, where the corneal architecture and density may be altered. Nevertheless, the data from this study suggest that the cell wall biosynthetic or regulatory proteins could serve as targets for prophylactic or early-intervention FK treatments. More broadly, this study highlights the fact that fungal virulence pathways are not universal and may vary markedly across host tissues due to salient differences in nutrient composition, immune cell dynamics, oxygen availability, or biomechanical properties. The altered gene expression and metabolic profiles of the fungus as a function of these variables will almost certainly impact antifungal sensitivity and, consequently, treatment outcomes. This of course complicates the development of novel antifungals, which not only must be tailored to specific fungal pathogens, but also to the affected tissue site.

## Acknowledgments

The authors would like to thank Mark Dittmar and staff at the Dean McGee Eye Institute (DMEI) Animal Research Facility and Louisa Williams at the DMEI Imaging Core for histology support.

---

**Supplementary Figure 1.**
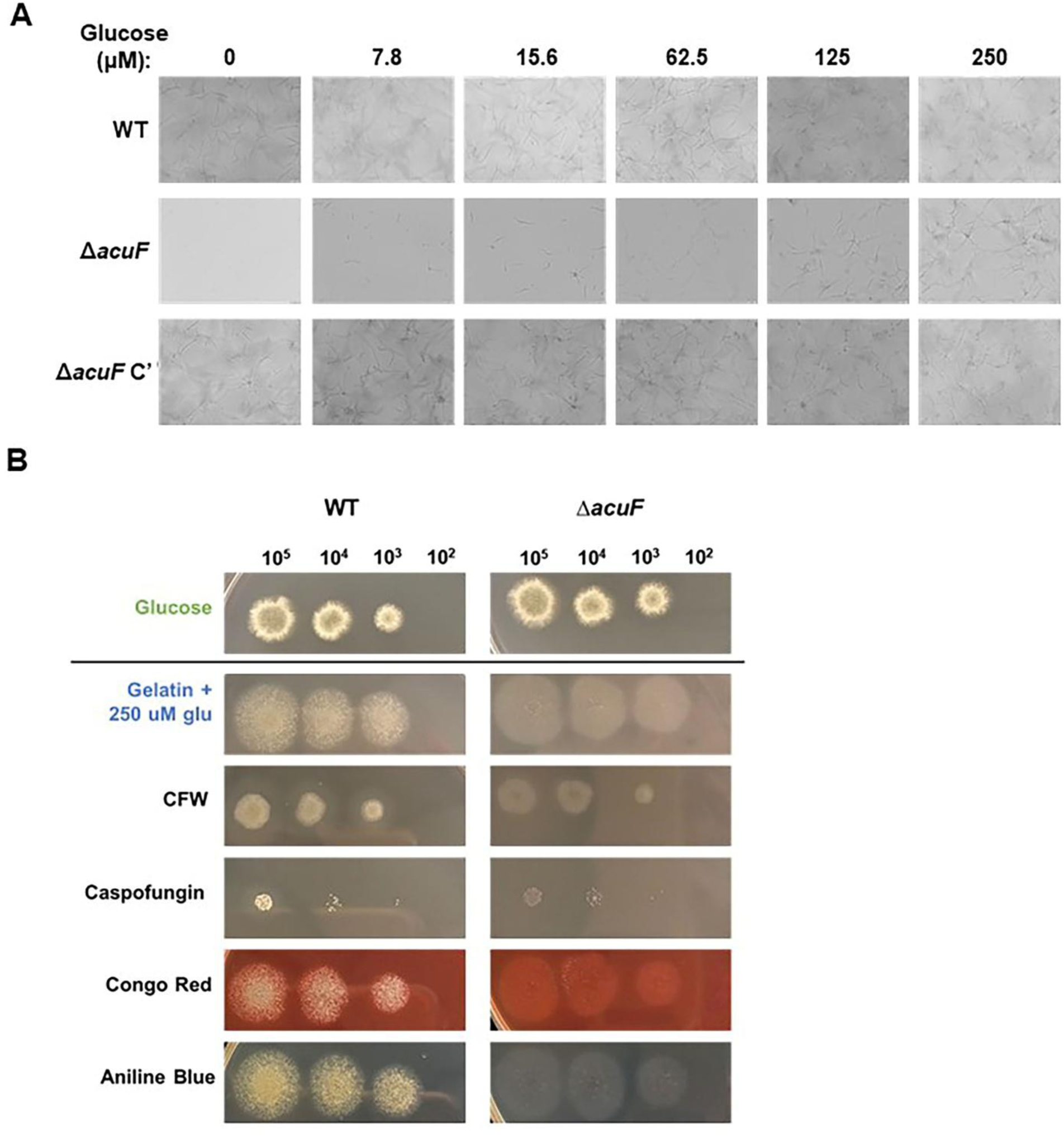
Glucose and cell wall stress sensitivity of *ΔacuF*. A) WT, Δ*acuF*, and Δ*acuF* C’ were inoculated at 10^5^ conidia/mL in acetate minimal media supplemented with various levels of glucose and incubated in 96 well plates at 35⁰C. Micrographs were taken at 48 h. **B)** Serial conidial dilutions (10^5^-10^2^) were spotted onto gelatin media supplemented with 250 μM glucose and the indicated cell wall stressors: calcofluor white (CFW), caspofungin, Congo red, or aniline blue. All compounds were added at 40 µg/mL.

**Supplementary Figure 2.**
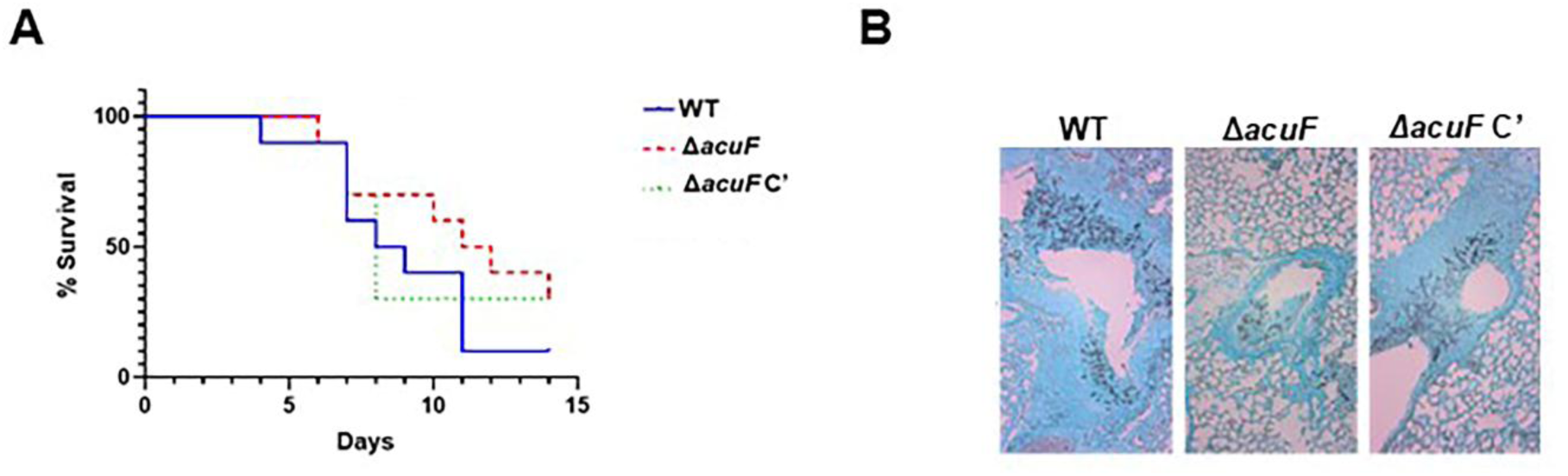
Loss of acuF does not impact disease establishment in a murine model of IPA. To assess pulmonary virulence, male CD-1 mice were immunosuppressed with triamcinolone and intranasally inoculated with 2×10^6^ conidia (WT, *ΔacuF*, or *ΔacuF C’*) or PBS as a control. **A)** The survival curves for each infection groups were indistinguishable by Logrank test (n=10/group); all groups were statistically different from a PBS (sham) inoculated control group (n=5). **B)** Grocott’s Methenamine Silver (GMS) staining of lung sections taken at 48 h p.i.

**Supplementary Figure 3.**
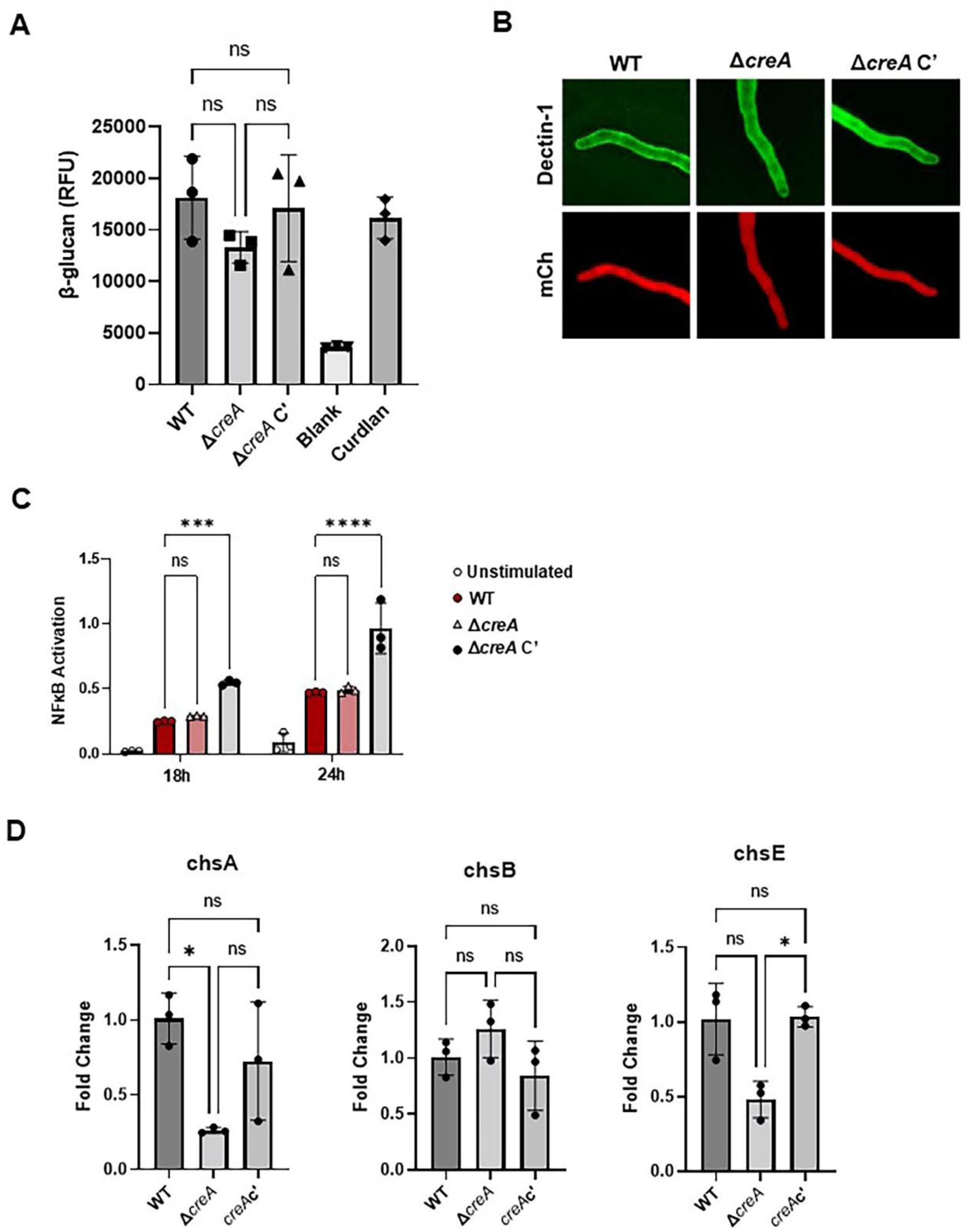
A) Strains were cultured for 48 h at 35⁰C and 250 rpm in liquid GMM AT. Fungal tissue was collected and dried overnight at 55°C. Dried tissue was then homogenized and solubilized in NaOH as described previously before being incubated in a 96-wells plate with aniline blue staining solution (0.067% aniline blue, 0.35N HCl, 0.98 M glycine-NaOH, pH 9.5) for 30 min at 50°C. Fluorescence in each well was measured at 405-nm excitation and 460-nm emission. **B)** WT, Δ*creA*, and Δ*creA* C’ were cultured for 18-22h in GMM, washed thrice with PBS, and fixed with 4% paraformaldehyde for 15 min at room temperature, followed by three more washes. Fixed hyphae were incubated for 2h with soluble Dectin-1 (1 µg/mL in PBS + 0.1% BSA, Invivogen, fc-mdec1a) at 4°C, washed with PBS+0.1% BSA three times, then incubated for 1h with FITC-conjugated rabbit anti-human IgG Fc (1:200 in PBS + 0.1% BSA, Thermofisher, 31535) at 4°C in the dark. Excess secondary antibody was removed by washing three more times with 0.1% BSA in PBS prior to imaging. Composite images were generated from z-stacks taken on an Olympus FV1200 confocal microscope. **C)** 1 × 10⁶ conidia were inoculated into 25 mL GMM in 250 mL Erlenmeyer flasks and incubated overnight at 37°C with shaking at 200 rpm. Hyphal suspensions were collected by centrifugation at 4500 rpm, washed 2–3 times with sterile phosphate-buffered saline (PBS), and disrupted by bead beating in PBS to generate hyphal fragments. Suspensions of hyphal fragments were then heat-inactivated at 65°C for 30 min, and fragment concentration was normalized across samples to an OD₃₆₀ = 0.3 in Dulbecco’s Modified Eagle Medium (DMEM). RAW-Blue™ cells (Invivogen) were seeded at 50,000 cells per well in 96-well flat-bottom plates in 100 μL complete DMEM (10% fetal bovine serum and antibiotics) and incubated overnight at 37 °C, 5% CO₂. The next day, 100 μL of heat-inactivated fungal suspension (OD₃₆₀ = 0.3) was added per well (final volume 200 μL). Control wells received 100 μL of plain DMEM. Supernatants were collected at 18 and 24 h post-stimulation. Secreted embryonic alkaline phosphatase (SEAP) activity in the supernatants was measured using Quanti-Blue™ reagent (InvivoGen) according to the manufacturer’s instructions. Groups were compared by Ordinary One-way ANOVA (****P<0.0001). **D)** Quantitative RT-PCR was performed on RNA isolated from WT, Δ*creA*, and Δ*creA* C’ cultured for 36 h in GMM. Statistics: Ordinary One-Way ANOVA (*p<0.05).

**Supplementary Figure 4.**
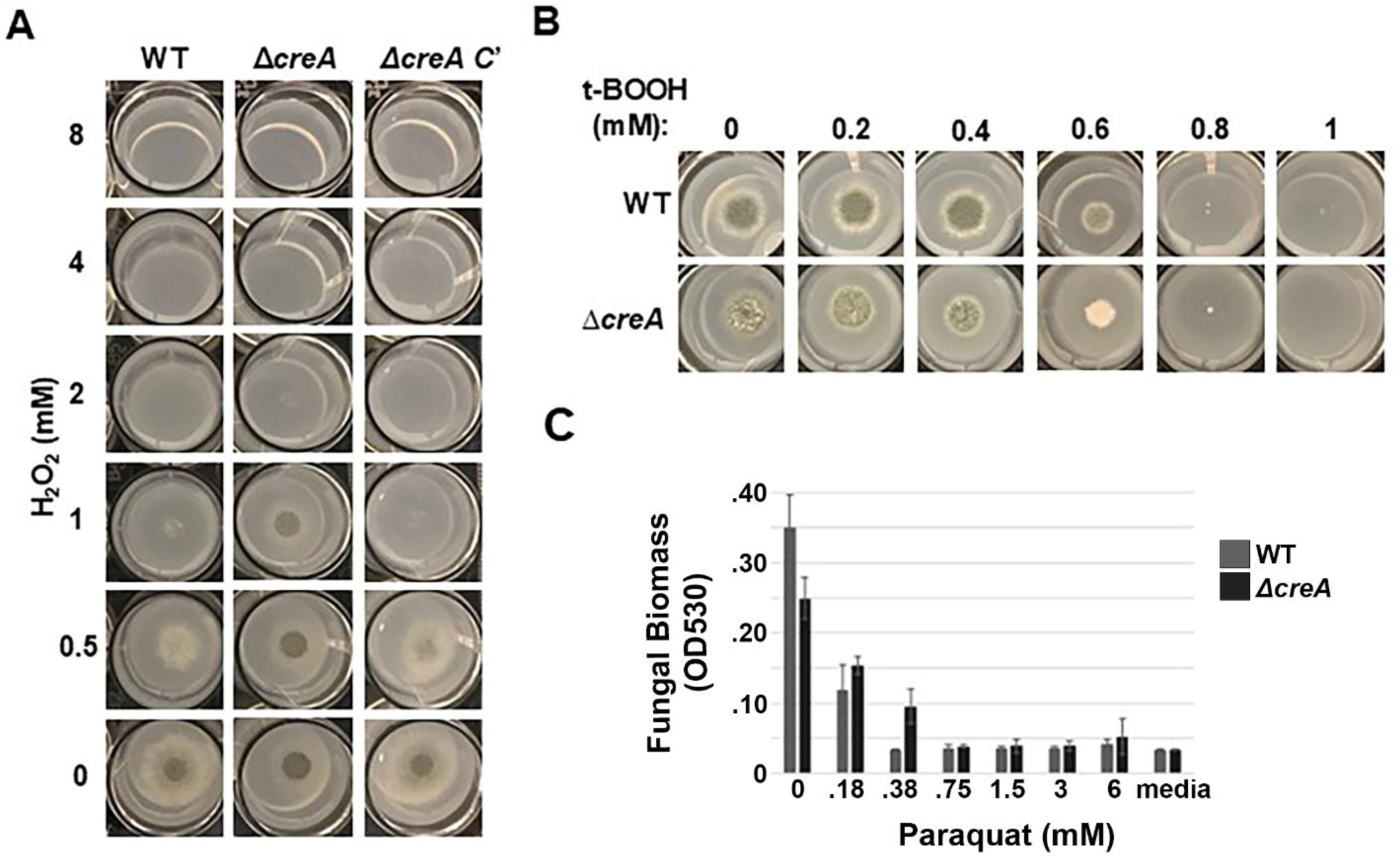
The *A. fumigatus creA* knockout is not hypersensitive to oxidative stressors. A-B) 2×10^3^ conidia were spotted onto GMM agarose containing the oxidative stressors H_2_O_2_ or t-BOOH at the indicated concentrations. Plates were incubated for 48 h at 35⁰C. **C)** Strains were inoculated into liquid GMM supplemented with 0-6 mM paraquat and incubated in a 96-well plate at 35⁰C for 48 h before growth was measured by taking the optical density of each well at 530 nm.

**Supplementary Figure 5.**
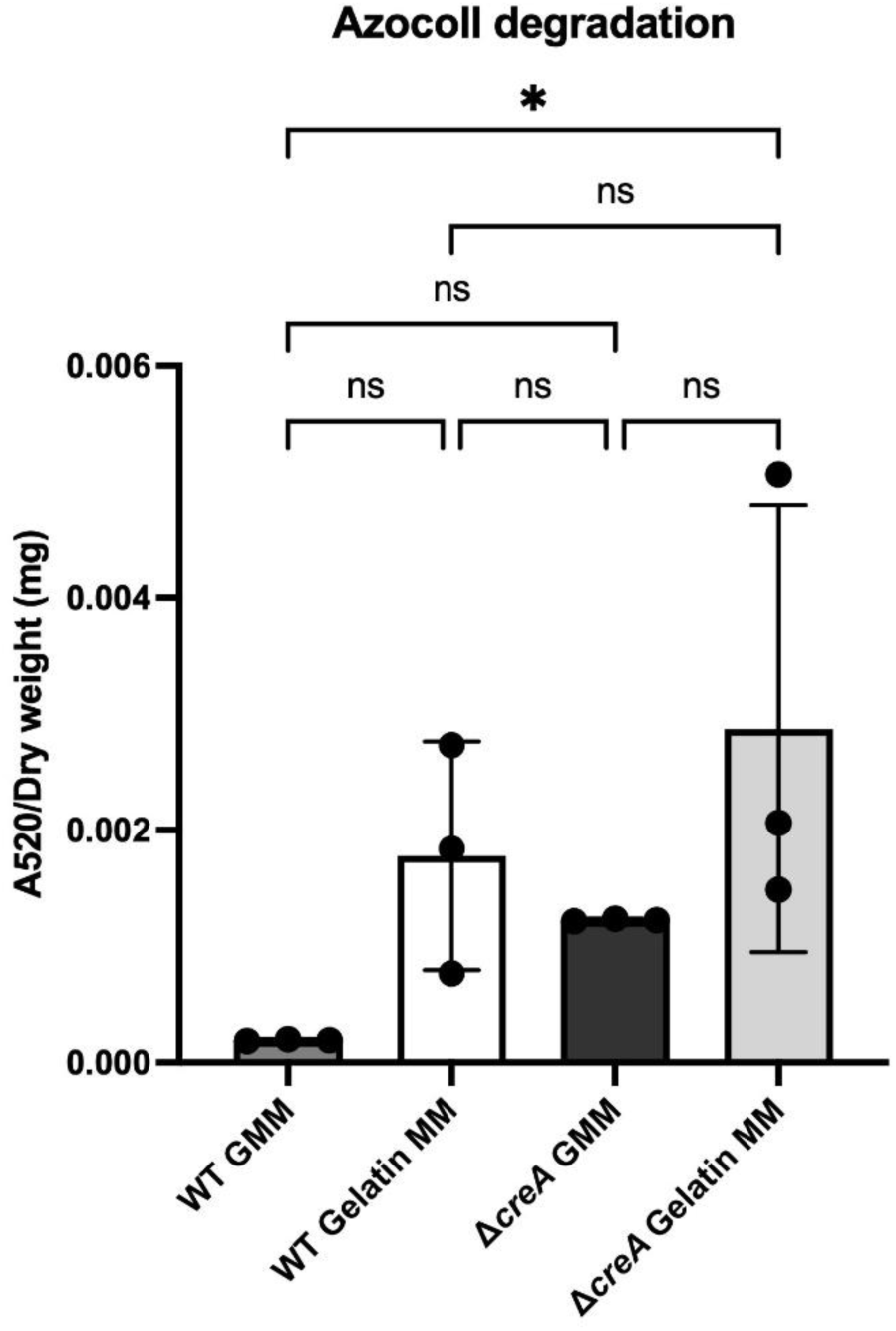
Extracellular collagenase activity is not altered in the *ΔcreA* mutant. 1 x 10^5^/mL conidia were inoculated into media containing 1% glucose (GMM) or 1% gelatin. Following incubation for 72 h at 35°C at (200 RPM, culture supernatants were incubated in the presence of pre-washed dye-labelled collagen (Azocoll (5 mg/mL) for 3 h at 37°C with gentle shaking. Subsequently, the tubes were centrifuged at 10,000 rpm for 3 min, followed by spectrophotometric measurement of absorbance at 520 nm to determine azo dye liberation from the supernatant. These values were normalized to the dry weight (g) of the individual strains using the 72 h biomass that was transferred to tubes and lyophilized for 48 h. Groups were samples analyzed by Kruskal-Wallis test * 0.0395.

